# Synergistic Combined-proteomics Guided Mapping strategy identifies mTOR mediated phosphorylation of LARP1 in nutrient responsiveness and dilated cardiomyopathy

**DOI:** 10.1101/2022.10.13.512080

**Authors:** Meng-Kwang Marcus Tan, Radoslaw M. Sobota, Esther SM Wong, Leah A. Vardy, Brian Burke, Colin L. Stewart

## Abstract

Increased activity of the mammalian target of rapamycin (mTOR) signalling pathway, a crucial nutrient sensor, exacerbates ageing and ageing-related diseases, including cancer and heart failure. To further elucidate the physiological role of the serine/threonine kinase mTOR, we devised a novel tractable proteomics strategy that combines interaction proteomics, proximity-based proteomics and quantitative phosphoproteomics to identify interactors with and potential substrates of mTOR. We identified 58 candidate mTOR substrates, several of which were further validated. Interestingly, several of these candidate mTOR substrates are involved in various aspects of RNA biology, including regulating stability and processing. We characterized in-depth one of the validated mTOR substrates, LARP1, an RNA binding protein. mTOR-dependent phosphorylation of LARP1 is nutrient-sensitive and controls the RNA-binding ability of LARP1. We show that mTOR activity and LARP1 and LARP1 phosphorylation levels are increased in a congenital mouse model of dilated cardiomyopathy (DCM) caused by a mutation in the Lamin A gene. This implicates LARP1 in the development of DCM.

## Introduction

The mTOR signalling pathway is an important orchestrator of cellular nutrient sensing and proliferation and is inhibited by rapamycin^1^. Central to this pathway is mTOR, a serine/threonine kinase, which is usually incorporated into two distinct kinase complexes; mTOR complex 1 (mTORC1) and mTOR complex 2 (mTORC2)^1^. mTORC1 primarily comprises the proteins RAPTOR, mLST8, PRAS40 and DEPTOR, while mTORC2 is made up of RICTOR, Protor-1/2, mLST8, mSin1 and DEPTOR. Besides having different protein components, mTORC1 and mTORC2 phosphorylate discrete substrates involved in many vital cellular processes, such as protein translation and autophagy^1^ that are primarily involved in the metabolic control of cell growth.

Due to its ability to regulate a broad and diverse array of cellular functions, misregulation of the mTOR pathway contributes to cancer progression and the development of metabolic and age-related diseases^2^. The connection between mTOR and ageing was established by the genetic and pharmacological ablation of mTOR signalling, in the nematode, *C. elegans*, fly, *Drosophila*, budding yeast, and mice, with these diverse organisms all showing enhanced longevity when mTOR was inhibited^3-8^. Besides extending lifespan, treating mice with the allosteric mTOR inhibitor, rapamycin and its derivatives, known as “rapalogs”, inhibited many ageing-related pathologies, including cancer and cardiac disease, specifically dilated cardiomyopathy (DCM)^9-14^. However, long-term or chronic usage of rapamycin results in glucose intolerance due to insulin resistance and immunosuppression, highlighting the need for alternative therapeutic treatments to regulate mTOR^15^. The key to the development of therapeutic strategies in reversing misregulated mTOR signalling in disease is identifying either regulators acting upstream of mTORC1 and mTORC2 or downstream effectors, especially downstream effectors mTORC1 and mTORC2 substrates. To this end, creative proteomics efforts have identified new upstream and downstream participants in the mTOR signalling pathway^16-21^. Though several mTOR substrates have recently been identified, the number of currently known mTOR substrates is insufficient to account for the far-reaching cellular effects of the mTOR signalling cascade. Hence, we sought to establish an innovative proteomics strategy to identify novel substrates of mTOR.

We devised the <u>S</u>ynergistic <u>C</u>ombined-proteomics <u>G</u>uided <u>M</u>apping (SCGM) strategy, which combines interaction-based proteomics, proximity-based proteomics and quantitative phosphoproteomics to aid in identifying mTOR substrates synergistically. The rationale for this strategy is that kinases phosphorylate their substrates by transiently interacting with them. The interaction-based proteomics aspect of the SCGM strategy involves affinity precipitation of a recombinant protein, which here is the mTOR protein, followed by mass spectrometry analysis^22-25^. This arm of the strategy allows for identifying proteins that either directly or indirectly interact with mTOR. In parallel, the proximity-based proteomics method, BioID, was performed by introducing an N-terminally biotin ligase, BioID2, fused–mTOR into HEK293T cells to identify proteins in close to or directly interacting with mTOR^26^. In addition, quantitative phosphoproteomics was performed through stable isotope labelling by amino acids in cell culture (SILAC) to determine the effects of pharmacological inhibition of mTOR signalling with the mTOR allosteric inhibitor, rapamycin, and mTOR catalytic inhibitor, Torin1, on the cellular phosphoproteome^20,27,28^. The SCGM strategy identified 58 mTOR-interacting proteins (excluding mTOR), involved in a host of cellular processes and whose phosphorylation status is sensitive to mTOR inhibition. With downstream validation steps and bioinformatics analyses, we determined that novel mTOR kinase substrates are present in this list of 58 proteins.

Among these novel mTOR kinase substrates is the RNA binding protein, LARP1, which binds to and regulates the stability and translation of a class of mRNAs containing a 5’-terminal oligopyrimidine sequence (5’ TOP mRNAs) and is overexpressed in various cancers^29-34^. We identified the phosphorylation of LARP1 at Serine 689 (S689) and Serine 697 (S697) as being mTOR signalling- and nutrient-sensitive. In addition, we determined that the phosphorylation of LARP1 at these sites is important in regulating the RNA binding ability of LARP1 to its target mRNAs, namely mRNAs encoding ribosomal proteins, such as RPS20 and RPL32^34,35^.

Among the diseases associated with elevated mTOR activity are laminopathies caused by mutations in the LaminA (*LMNA*) gene. Patients and mice with a mutation in *LMNA* where the amino acid asparagine at position 195 is replaced with a lysine (*Lmna*^N195K^) develop DCM, a heart disease characterized left ventricular chamber dilation and wall thinning^36,37^. In the *Lmna*^N195K^ mouse model, we find that mTOR signalling and the phosphorylation of LARP1 are elevated in the hearts of mice with DCM. The identification and characterization of the mTOR-dependent phosphorylation of LARP1 not only provide an insight into the regulation of LARP1 function it further highlights the robustness of the SCGM strategy in the identification of mTOR substrates. Moreover, we identified LARP1 as a potentially novel factor contributing to the development of DCM.

## Results

### Identifying phosphoproteome changes induced by inhibiting mTOR kinase

The mTOR signalling pathway, being a nutrient-sensing pathway, responds to nutrient fluctuations^38^. We first tested the effects of starving mouse adult fibroblasts (MAFs) by culture in Hanks’ Balanced Salt Solution (HBSS) for 1 hour, followed by 3 hours incubation with complete media with (i.e. with Foetal Calf Serum), without mTOR inhibitors of the mTOR signalling pathway (Supplementary Figure 1A). Our results showed that MAFs starved with HBSS for an hour have significantly reduced levels of phosphorylated EIF4EBP1, a well-characterized mTOR kinase substrate^39^. When the medium in which the MAFs were maintained was replaced with complete media, mTOR signalling was restored, as determined by an increase in the levels of phospho-EIF4EBP1 to the levels seen in the non-HBSS incubated MAFs. In contrast, phospho-EIF4EBP1 levels in MAFs incubated with complete media with the mTOR inhibitor Torin1 were not increased (Supplementary Figure 1B).

We initiated our phosphoproteomics analysis by starving the SILAC labelled MAFs of nutrients for 1 hour. After an hour, complete media and the corresponding inhibitors were added to the three sets of SILAC-labelled MAFs. The controls were treated with DMSO, while one set was treated with the mTOR allosteric inhibitor, rapamycin, and the second group was treated with mTOR catalytic inhibitor, Torin1. All three groups of MAFs were incubated for a further 3 hours (referred to as “Forward Labelling” in Figure 1A). To account for potential phosphoproteome changes that might be due to differential SILAC labelling across the three groups of MAFs, the same experiment was also performed, but this time, the compound treatment to each group of the SILAC labelled cells were switched (refer to “Reverse Labelling” in Figure 1A for more details). For each SILAC experiment, equal numbers of cells from each group of MAFs were mixed and lysed. The proteins in the lysates were proteolytically cleaved with Lysyl Endopeptidase, LysC, and trypsin. The resultant peptides were fractionated by strong cation exchange (SCX) into different fractions. The phosphopeptides were enriched from the fractions by titanium dioxide treatment. The identities of the phosphopeptides and the quantitative differences in the phosphoproteomes in the MAFs across the three treatments were analyzed by mass spectrometry (MS) (Figure 1A). For both the Forward and Reverse Labelling experiments, the ratios of peptides found in the compound-treated sample to similar peptides found in the control DMSO-treated sample were first normalized by multiplying the raw ratio with a factor obtained, taking into account differences in protein abundance between the two samples (Supplemental Table S1). Proteins with peptides with two or more fold changes in the compound-treated samples in both the Forward and Reverse Labelling experiments were considered to be regulated by mTOR kinase inhibition (Supplemental Table S2).

**Figure 1:**
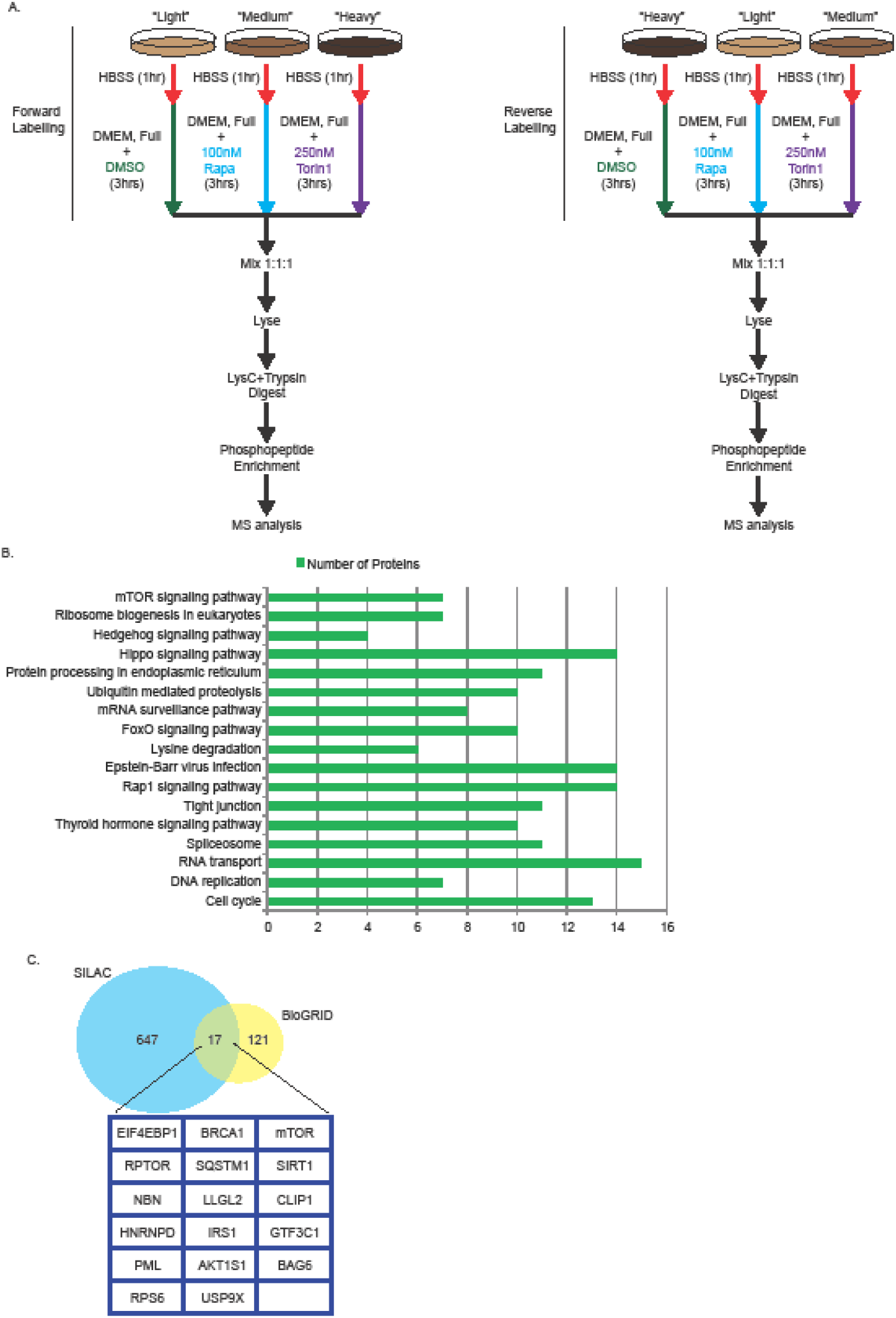
Summary of SILAC based phosphoproteomics procedure and results. A) Flow chart depicting the key steps involved in the SILAC based phosphoproteomics. B) Gene Ontology (GO) analysis (using default parameters) performed on the proteins whose phosphorylation levels were reduced upon mTOR inhibition to determine the biological processes regulated by these proteins (p<0.05). C). Cross-referencing proteins whose phosphorylation levels decreased after mTOR inhibition was identified using the SILAC based phosphoproteomics approach with known mTOR interactions curated in the protein interaction database, BioGRID.

The identified proteins were analyzed using the Database for the Annotation, Visualization and Integrated Discovery (DAVID) and classified according to their Gene Ontology (GO) terms to determine the significant pathways affected by mTOR inhibition^40-42^. This analysis identified one of the affected pathways as the mTOR signalling pathway (p<0.05). Other pathways affected by mTOR inhibition include ribosome biogenesis, hedgehog signalling, hippo signalling and the ubiquitin-mediated proteolysis pathways, all linked to mTOR signalling (Figure 1B). Besides identifying cellular pathways affected by mTOR inhibition, we determined the number of known mTOR interactors from our protein list obtained from the SILAC approach. To do so, we compared the proteins identified from our SILAC approach with a list of known mTOR interactors curated in the Biological General Repository for Interaction Datasets (BioGRID)^43^. From this analysis, 17 known interactors of mTOR displayed a reduction in phosphorylation levels after mTOR inhibition, and among these 17 proteins were known mTOR substrates such as EIF4EBP1 and IRS1 (Figure 1C). These results confirmed that our drug treatment protocol and subsequent phosphoproteomics analysis is a valid approach to identify proteins whose phosphorylation is regulated by mTOR.

### Identification of candidate mTOR substrates from data obtained by SILAC

Concurrent with using SILAC, we performed two other proteomics analyses to identify proteins, among the list identified from SILAC, that are candidate substrates of mTOR. The mTOR proteins in human, mouse and rat share a high degree of sequence homol-ogy (data not shown). Hence, human and rat mTOR proteins were used in the interac-tion proteomics to validate the interactors identified. One of the proteomics procedures selected is interaction proteomics using immunoprecipitation of MYC-BioID2 tagged mTOR (from human and rat) followed by mass spectrometry (IP-MS) analysis (Figure 2A). This procedure was chosen as it identifies proteins that form complexes with mTOR. To do this, the interactors identified in the human and rat mTOR IP-MS analyses displayed a two-fold increase in abundance (after normalization with the bait levels) over those identified in the EGFP IP-MS sample were regarded as candidate interactors of mTOR (Supplemental Table 3). After comparing known mTOR interactors obtained from BioGRID, we identified many known mTOR interactors, including mTOR substrates, in our IP-MS data (Supplemental Figure 2). While the mTOR IP-MS procedure identifies proteins that complex with mTOR, it may not allow for the recovery of potential mTOR substrates due to the transient nature of enzyme-substrate interactions^44^. To complement the IP-MS approach, we utilized the proximity-based proteomics BioID procedure to search for mTOR interactors and substrates. With BioID, proteins in proximity (10-20nm) to mTOR are labelled with Biotin by BioID2, a biotin ligase derived from *Aquifex aeolicus*, which is fused to the amino-terminus of mTOR^45^. The biotinylated peptides are isolated using Streptavidin antibody-conjugated beads (Figure 2B). This proximity-based proteomics approach identifies potential substrates that may no longer be complexed with mTOR but are still close to mTOR before or after phosphorylation by mTOR. This procedure identified many known mTOR interactors (including mTOR substrates) after comparing the BioID data with BioGRID-curated mTOR interactors (Supplemental Figure 2 and Supplemental Table 4).

**Figure 2:**
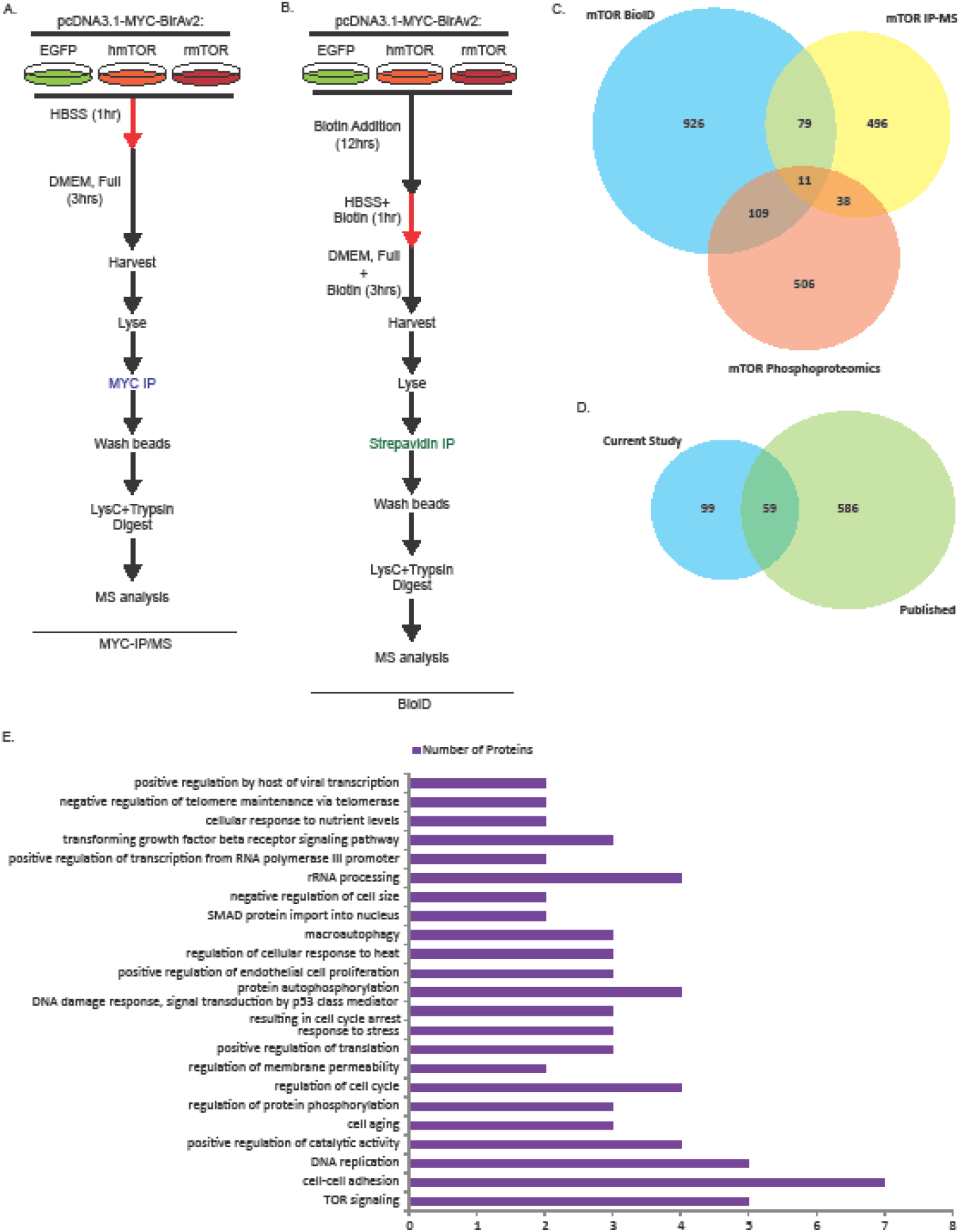
Identification and classification of high confidence novel candidate substrates of mTOR. A & B) Flow chart depicting the key steps involved in the mTOR-interaction proteomics and BioID approaches. C) Venn diagram illustrates the identification of proteins interacting or in close proximity to mTOR and whose phosphorylation levels are mTOR signalling dependent using the SCGM strategy. D) Identifying high confidence candidate mTOR substrates by cross-referencing proteins identified using the SCGM strategy with data from three published mTOR phosphoproteomics studies. E) Gene Ontology (GO) analysis (using default parameters) performed on the high confidence candidate mTOR substrates (p<0.05).

To identify potential mTOR interactors, we applied our Synergistic Combined-proteomics Guided Mapping (SCGM) strategy by combining and comparing data from all 3 of our proteomics approaches. Through this strategy, 158 proteins (including mTOR) are found either to interact with mTOR or are close to mTOR, and whose phosphorylation status is sensitive to mTOR inhibition (Figure 2C). Through different methods of perturbing the mTOR signalling pathway, three recent phosphoproteomics studies identified protein phosphorylation events dependent on mTOR signalling^18-20^. However, the data from these studies do not distinguish between proteins that are substrates of mTOR and proteins which are indirectly phosphorylated by mTOR-regulated kinases, such as S6K1 and AKT1^1^. Therefore, to further increase the probability of identifying mTOR substrates, we compared the identities of our 158 candidate proteins with those found in the previous three studies and identified 59 (including mTOR) proteins that are common across all four studies (Figure 2D). Bioinformatics analysis of the 59 candidate proteins revealed these proteins participate in numerous pathways, including the mTOR, ErbB and AMPK signalling pathways, regulating a diverse range of biological processes, including nutrient response and autophagy DNA damage and cell ageing (Figures 2E and Supplemental Figure 3).

### Validation of candidate proteins obtained through SCGM

From the 58 candidates (excluding mTOR), we next determined if these candidate proteins do indeed bind to mTOR and are sensitive to mTOR. First, we selected six proteins and tested them to see if these proteins responded to mTOR inhibition. We treated either regular HEK293T cells or HEK293T cells transiently expressing the hemagglutinin (HA)-tagged protein-of-interest with either DMSO or 250nM Torin1 4 hours. After 4 hours, the cells were harvested, and the lysates used for immunoblotting by SDS-PAGE gels containing Phos-Tag^™,^ which separates phosphorylated forms of a protein from its non-phosphorylated counterpart. As a positive control, for each of the six proteins-of-interest, the lysate for DMSO treated sample was split into two, with one incubated with Lambda phosphatase (λPPase) for 1 hour to remove all phosphorylated residues on the protein-of-interest. As demonstrated in the blot of a well-known mTOR substrate, EIF4EBP1, Torin1 treatment reduced the amount of slower migrating phosphorylated forms of EIF4EBP1 and increased the faster migrating non-phosphorylated EIF4EBP1. In addition, λPPase-treatment significantly reduced phosphorylated forms of EIF4EBP1 (Figure 3A). A similar trend was observed for five of the proteins of interest, indicating that the phosphorylation levels on these proteins are reduced by mTOR inhibition.

**Figure 3:**
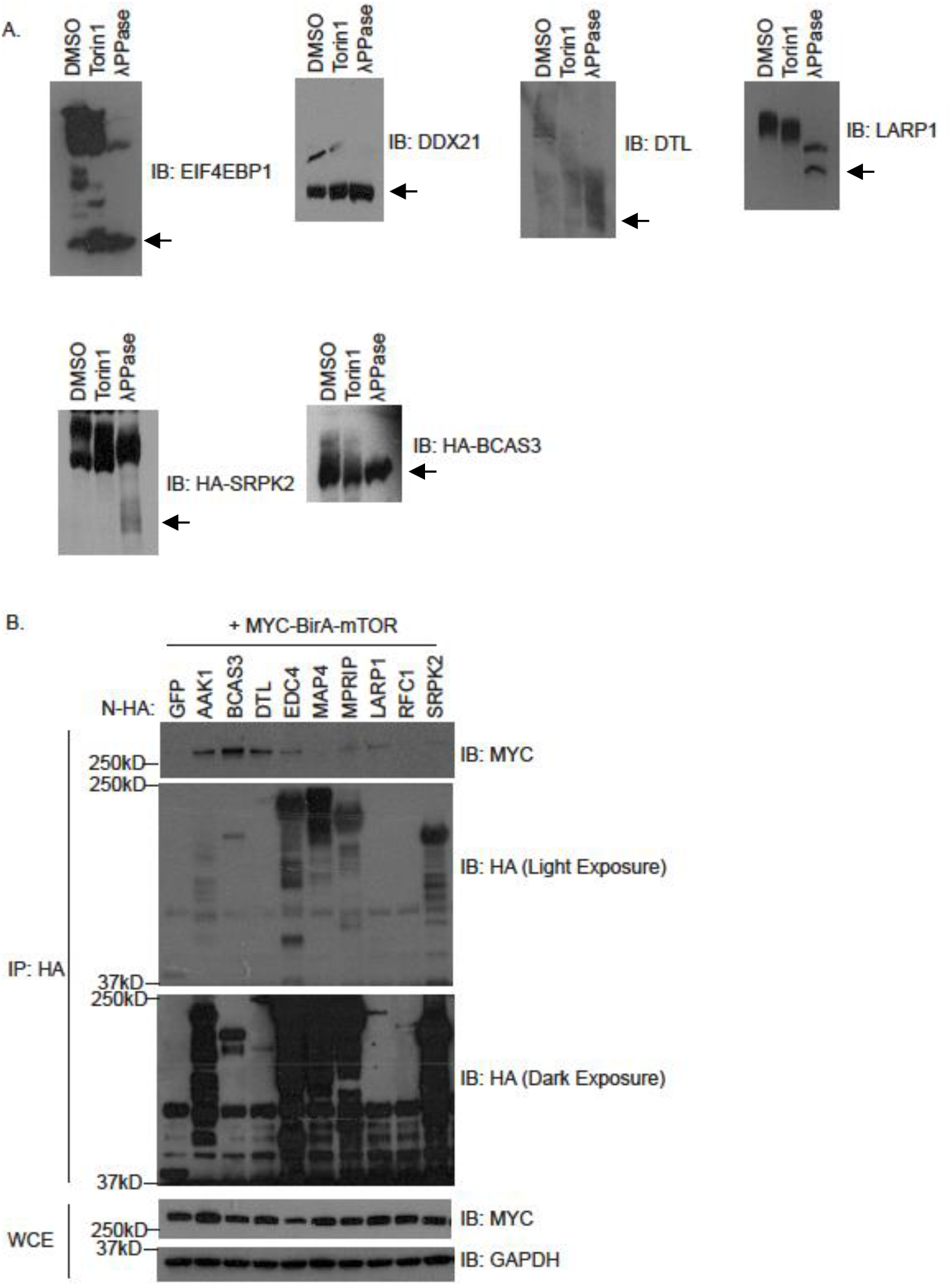
Validation of high confidence novel candidate mTOR substrates. A) Blots of lysates of cells treated 4 hours with either DMSO or 250nM Torin1. For each blot, the last lane represents lysate that was treated with Lambda Protein Phosphatase for an hour. The samples were run on a PhosTag^™^ gel followed by subsequent immunoblotting analysis with the indicated antibodies. (← indicates non-phosphorylated form of protein) B) Immunoprecipitation of HA-tagged EGFP or indicated candidate substrate from transfected cells to determine their binding to MYC-BioID2-mTOR.

In addition, we selected nine candidate proteins, fused to an HA-tag at each of their N-terminus and transiently co-expressed each of these proteins with MYC-BioID2 tagged mTOR in HEK293T cells (Figure 3B). Immunoprecipitation with HA-antibody conjugated beads followed by immunoblotting was performed to determine the ability of each of these nine proteins to bind to MYC-BioID2 tagged mTOR. Our results indicated that 7 out of the nine selected proteins interact with mTOR (Figure 3B). Combining the two validation approaches, DTL, LARP1, SRPK2 and BCAS3 displayed the two hallmarks of a typical kinase substrate, namely, 1) the substrate interacts with the kinase and 2) perturbations to the kinase affect the phosphorylation of the substrate, suggesting that there is a high likelihood that they are mTOR substrates (Figures 3A and 3B). LARP1 interacts with mTOR, and at the peptide level, two phosphopeptides from LARP1 are sensitive to mTOR inhibition, strongly suggesting that LARP1 is an mTOR substrate but the biological consequences of mTOR phosphorylation of LARP1 are unknown^34,35,46^.

Certain kinases have preferences for specific amino acid motifs found on their substrates, and mTOR has a strong preference for proline-, phenylalanine-, leucine-directed serine/threonine motifs on its target proteins^20^. We mapped potential mTOR target sites on the 58 candidate proteins based on the presence of proline-, phenylalanine-, leucine-directed serine/threonine found on peptides that showed changes in phosphorylation levels seen in our SILAC approach (Figure 4). We began this analysis by identifying peptides with changes in phosphorylated serine and/or threonine residues in the Forward, and Reverse SILAC approaches. After identifying the positions of these serine and threonine residues on their respective proteins, we compared our results with the PhosphoSitePlus®, a repository for protein modification, to identify sites previously reported as being phosphorylated. Phosphorylated serine and threonine residues were then examined to determine if a proline, phenylalanine or leucine residues were present immediately after the phosphorylated serine/threonine (Supplemental Table 4)^47^. As presented in Figure 4, most of the 58 candidate proteins have serine/threonine residues found in motifs targeted by mTOR. However, the physiological consequences of the phosphorylation of many of these residues by mTOR are still unclear.

**Figure 4:**
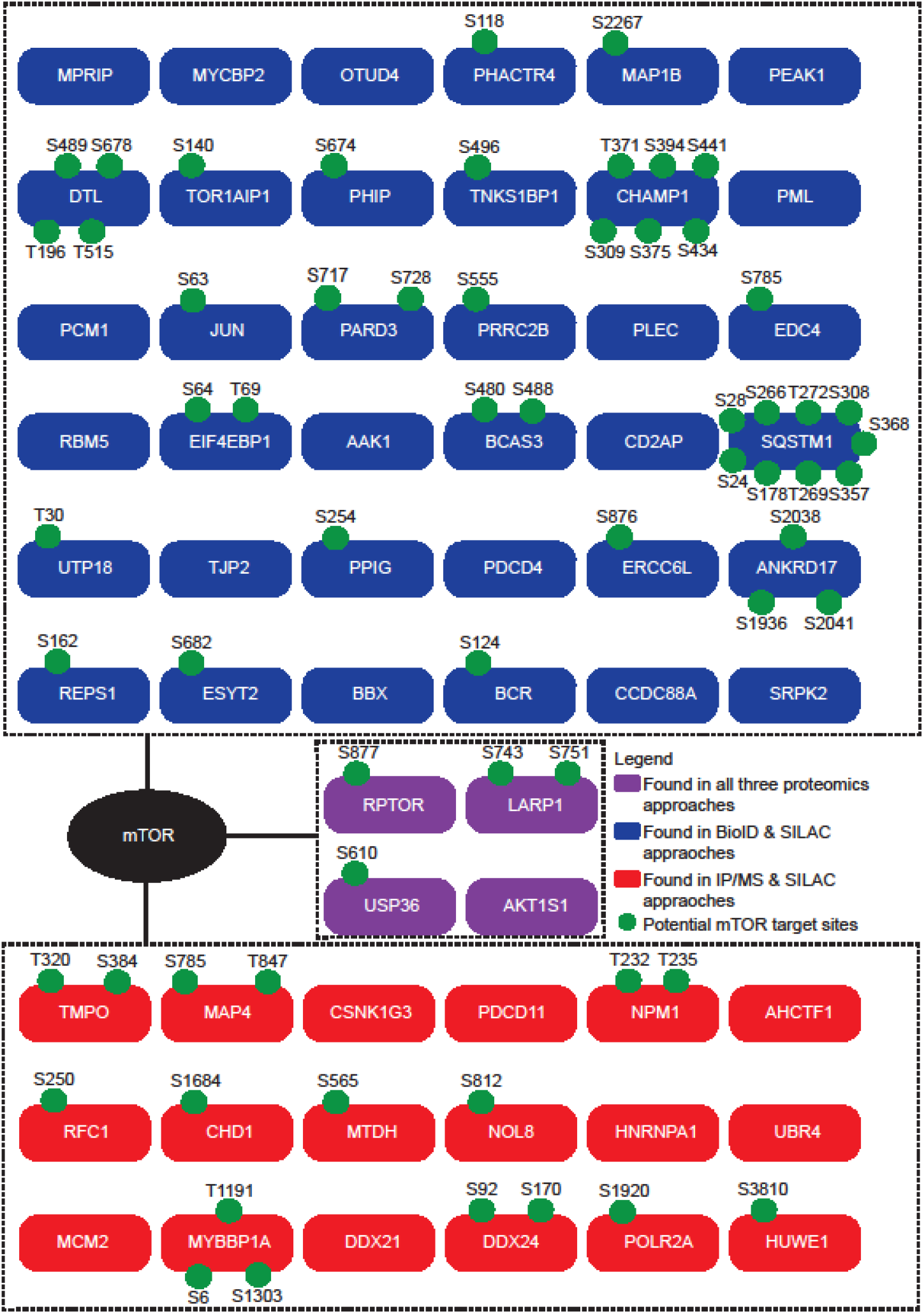
Identities of high confidence novel candidate mTOR substrates and location of possible mTOR target sites identified by the SCGM strategy. Fifty-eight high confidence novel candidate mTOR substrates (excluding mTOR) were grouped according to the proteomics approaches responsible for determining them. In purple are proteins found in all three proteomics approaches; in blue are proteins found in the BioID and SILAC-based proteomics approaches; in red are proteins found in the interaction proteomics SILAC-based proteomics approaches. Green circles represent possible mTOR target sites on high confidence novel candidate mTOR substrates. These potential mTOR target sites (found in the mouse homolog of the listed proteins) were selected based on three criteria: first, the phosphorylation on these target sites was found to have changed by two-fold in both the Forward and Reverse SILAC-based proteomics experiments; second, these target sites are mainly proline-, phenylalanine or leucine-directed – similar to known mTOR target sites; third, these target sites have been documented and curated on PhosphoSitePlus®^47^. For ease of reference, protein identities are based on the human homologs of the candidate proteins (found in mouse) listed in Supplemental Table 4.

### mTOR interacts with and phosphorylates LARP1 at S689 and S697

Our results validated the interaction between ectopically co-expressed mTOR and LARP1 (Figure 3B). To determine if the mTOR-LARP1 interaction is physiological, we ectopically expressed either mTOR/LARP1, performed an immunoprecipitation to pull down the ectopically expressed protein followed by immunoblotting to determine its interaction with its endogenous counterpart (Figures 5A and 5B). As an alternative, we immunoprecipitated endogenous mTOR/LARP1 and looked for its interaction with its endogenous partner (Figure 5C). Through these methods, we confirmed mTOR’s interaction with LARP1. In addition, we showed by immunofluorescence that RPTOR, a key component of the mTORC1 complex, colocalizes with LARP1 in the cytoplasm (Figure 5D). Our SILAC and motif analyses determined that Serine 743 and Serine 751 on mouse LARP1 are potential mTOR target sites (Figure 4), confirming a previous study showing that peptides of LARP1 containing these residues are phosphorylated by mTOR^46^. We then determined if phosphorylation of LARP1 by mTOR occurs at the protein level. A homology analysis of the LARP1 protein showed that the amino acid residues in the region around Serine 743 and Serine 751 are highly conserved across species, including mice, humans and zebrafish (Figure 5E). Serine 743 and Serine 751 in mouse LARP1 corresponds to Serine 689 and Serine 697 in full-length human LARP1. To study phosphorylation of S689 and S697 in human LARP1, we generated antibodies targeting the phosphorylation of these sites. These antibodies were then tested to determine if they specifically recognize their corresponding target phosphorylation sites and their efficacy in mouse samples. We confirmed that the antibodies targeting phosphorylated S689 (α-pS689) and phosphorylated S697 (α-pS697) specifically recognize their corresponding target phosphorylation sites (Supplemental Figure 4A). In terms of usage on mouse samples, only α-pS689 managed to detect a change in band intensity in MAFs treated with/without mTOR inhibitor, Torin1 (data not shown).

**Figure 5:**
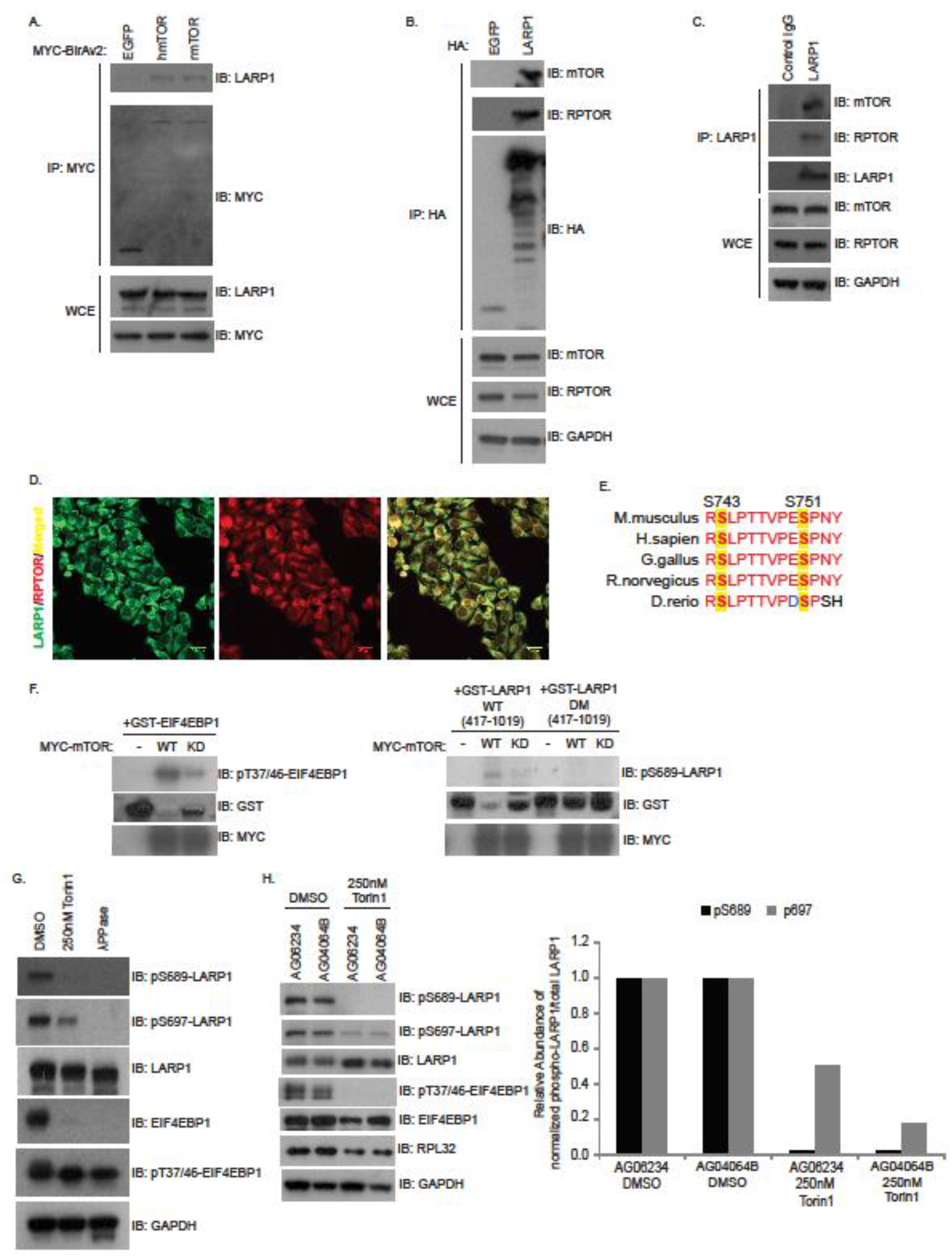
mTOR phosphorylates LARP1. A) Immunoprecipitation of MYC-BioID2-tagged EGFP, human mTOR (hmTOR) or rat mTOR (rmTOR) and subsequent immunoblotting with indicated antibodies. B, C) Immunoprecipitation of either ectopically expressed HA-EGFP or HA-LARP1 (B), or endogenous LARP1 (C), followed by immunoblotting with indicated antibodies. Figures A, B, C shows that mTOR and LARP1 interact with each other. D) Immunofluorescence with the indicated antibodies demonstrates the colocalization of LARP1 with mTORC1-component, Raptor. E) Comparison of mTOR target sites on LARP1 found in MAFs across different species shows high amino acid sequence conservation around these sites across different species. F) In vitro kinase using bacterial purified GST-EIF4EBP1, truncated wild type LARP1 [GST -LARP1_WT (417-1019)] and truncated LARP1 double mutant with mTOR target sites at S766 and S774 mutated to alanines [GST -LARP1_DM (417-1019)] as substrates and wild type (WT) and kinase-dead (KD) mTOR as the kinase. HEK293T cells (G) and human primary fibroblasts (AG06234 and AG04064B) (H) were treated with either DMSO or 250nM Torin1 for 4 hours. After 4 hours, the cells were lysed, and the lysates were used for subsequent immunoblotting analysis using the appropriate antibodies. In the last lane in (G), half the lysate obtained from the DMSO treated sample was incubated with Lambda Phosphatase to reduce the levels of phosphorylated proteins.

Consequently, in experiments involving mouse samples, only α-pS689 was used to detect changes in LARP1 phosphorylation. With these phospho-specific antibodies, we then determined if LARP1 is phosphorylated by mTOR *in vitro*. After incubating bacterial purified GST tagged truncated wild type LARP1 (GST-LARP1 WT) and double mutant LARP1 (GST-LARP1 DM), which has both S689 and S697 substituted with alanines, with either wild type (WT) or kinase-dead (KD) mTOR in the presence of ATP, the samples were analyzed by immunoblotting to determine the degree of LARP1 phosphorylation in each of the samples. The phosphorylation of GST-LARP1 WT increased significantly in the presence of WT mTOR but not in the company of KD mTOR, similar to that seen in the positive control GST-EIF4EBP1 (Figure 5F). Subsequently, we asked if mTOR inhibition affects LARP1 phosphorylation. In both an immortalized cell line (HEK293T) and primary human fibroblasts, inhibition of mTOR activity by mTOR catalytic inhibitor, Torin1, significantly diminished the levels of phosphorylated S689and S697 residues in LARP1 (Figures 5G and 5H). Our results establish LARP1 as a substrate of mTOR.

### Physiological and disease relevance of LARP1 phosphorylation

By confirming the phosphorylation of human LARP1 on S689 and S697 by mTOR, we determined the physiological relevance of the phosphorylation of LARP1 at these sites. As mTOR signalling is nutrient-sensitive, we determined that if LARP1 phosphorylation is regulated by mTOR signalling, conditions restricting mTOR signalling should also reduce LARP1 phosphorylation. To test this, we subjected HEK293T cells to 2 conditions that inhibit mTOR signalling, namely, nutrient starvation and pharmacological inhibition of mTOR using Torin1. HEK293T cells starved of nutrients displayed a dramatic reduction in LARP1 phosphorylation as seen from the decrease in the levels of phosphorylated S689, demonstrating nutrient sensitivity of LARP1 phosphorylation (Figure 6A). The same decrease in phosphorylated S689 was also observed in cells treated with Torin1 (Figure 6A). The ability of LARP1 to respond to nutrient abundance led us to determine if LARP1 responded to other cellular stresses. To do so, we treated C2C12 cells with a panel of compounds to induce various cell stresses, including hypoxia and proteotoxic stress. While serum starvation and proteasome inhibition by Bortezomib decreased phosphorylated LARP1, cytotoxic stress brought about by MLN4924 treatment, a neddylation inhibitor known to trigger cytotoxic stress responses, increased the levels of phosphorylated LARP1 (Figure 6B)^48,49^. Interestingly, this panel of stressors triggers the phosphorylation of EIF4EBP1 like the phosphorylation of LARP1 (Figure 6B).

**Figure 6:**
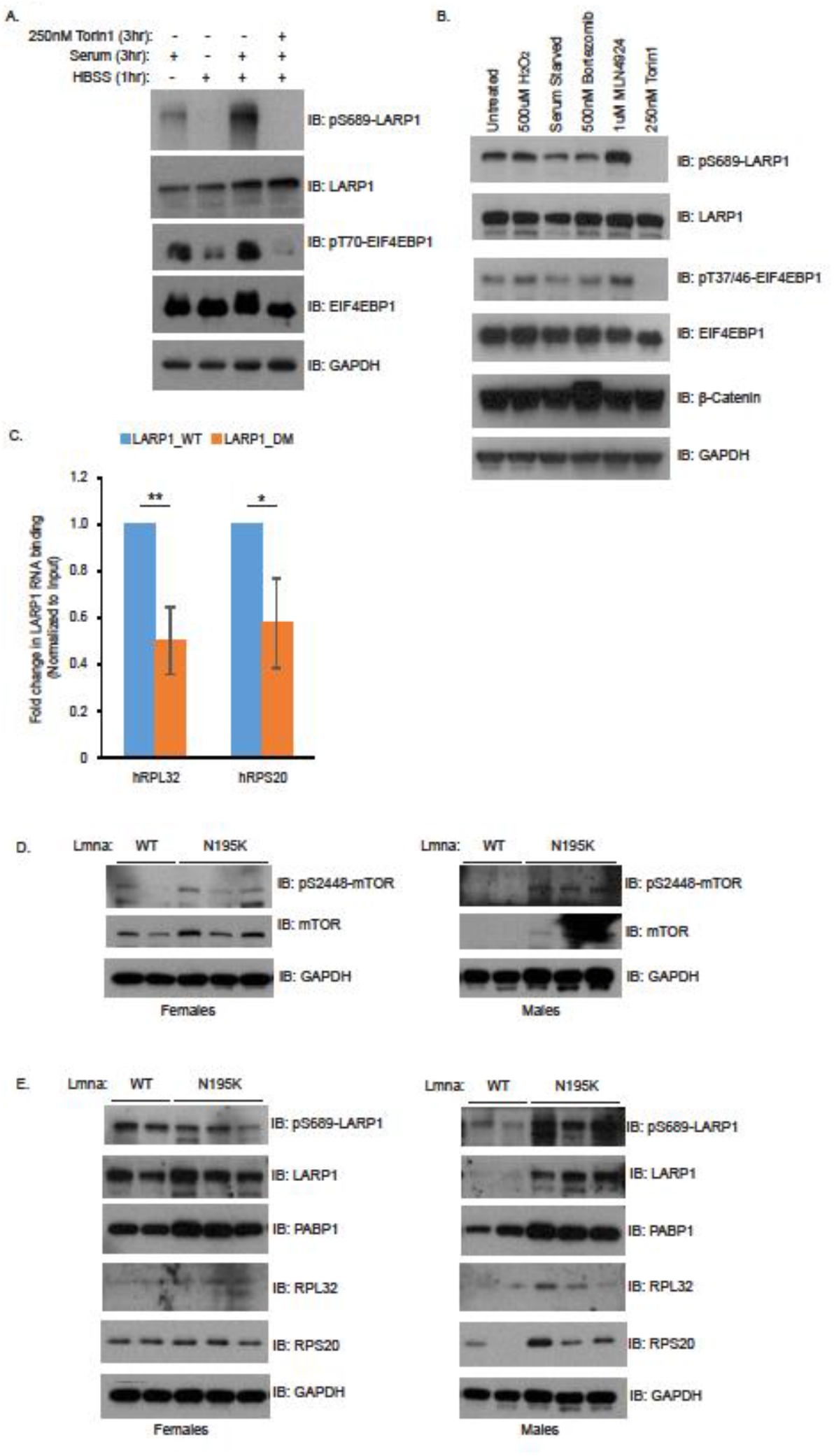
Phosphorylation of LARP1 is sensitive to mTOR inhibition, affects LARP1 RNA binding ability and is linked to DCM. A) HEK293T cells were subjected to the indicated treatments, and the lysates obtained from subsequent lysis were analyzed by immunoblotting with the appropriate antibodies. B) HEK293T cells were subjected to the indicated treatments, and the lysates obtained from subsequent lysis were analyzed by immunoblotting with the appropriate antibodies. C) HEK293T cells expressing transiently transfected HA-LARP1-WT or HA-LARP1-DM were lysed, and the lysates were subjected to RNA immunoprecipitation with agarose beads conjugated with HA-antibody. After immunoprecipitation, samples were processed, and the amount of RPL32 and RPS20 transcripts bound to HA-LARP1-WT and HA-LARP1-DM were assessed by qPCR. **p<0.005, *p<0.05, mean±SEM (n=3). Statistical analysis (unpaired *t*-test) was performed using the Prism software. D, E) Hearts from 9-week old male and female wild-type (WT) or Lmna N195K mutant (N195K) mice were harvested and analyzed by immunoblotting with the appropriate antibodies to compare the levels of mTOR signalling (D) and LARP1 phosphorylation (E) in these mice.

Given that LARP1 binds to components of the protein translation machinery and regulates the stability of 5’ TOP containing mRNAs, which encodes many ribosomal proteins and components of the translation machinery, we determined how the phosphorylation of LARP1 by mTOR might affect these properties of LARP1^33-35^. LARP1 preferentially binds to specific 5’ TOP containing ribosomal mRNAs, and two of these are transcribed from the RPS20 and RPL32 genes^34^. We then determined if phosphorylation of LARP1 by mTOR affects the ability of LARP1 to interact with RPS20 and RPL32 transcripts. Through RNA immunoprecipitation, we found that removing the mTOR phosphorylation sites on LARP1 reduced the interaction between LARP1 and RPL32/RPS20 transcripts (Figure 6C). In addition to lowering LARP1-RPL32/RPS20 mRNA interaction, mTOR inhibition by Torin1 also led to a reduction in the levels of RPL32 (Figure 5G).

While many studies have implicated LARP1 in cancer progression, the relevance of LARP1 in other diseases is still unknown^29-31,50^. Recently, several studies have established a link between mTOR signalling and DCM, characterized by left ventricular enlargement of the heart^13,14^. In two mouse models of DCM, where the *Lmna* gene in these mice was either deleted or carried a missense mutation (H222P), mTOR signalling was elevated compared to normal mice^13,14^. Since we identified and verified LARP1 as a mTOR substrate, we used another mouse model of DCM, the *Lmna*^N195K^ line, to determine if mTOR signalling was also elevated in this DCM model and how changes in mTOR signalling affect LARP1 phosphorylation. Intriguingly, we observed that mTOR signalling, as indicated by the levels of phosphorylated EIF4EBP1, is elevated in male *Lmna*^N195K^ mice but not in females (Figure 6D), which correlates with *Lmna*^N195K^ females living longer than *Lmna*^N195K^ males. Together with the increase in mTOR signalling, there is an increase in the levels of LARP1 and phosphorylated LARP1 in male *Lmna*^N195K^ mice (Figure 6E). Besides elevated mTOR signalling and phosphorylated LARP1, increases in RPS20 and RPL32 protein levels were also observed in male *Lmna*^N195K^ mice. Taken together, our data suggest that mTOR-dependent phosphorylation of LARP1 is nutrient-sensitive and regulates the binding of LARP1 to its target transcripts. Furthermore, our data establish a link between LARP1 and the development of DCM.

## Discussion

The mTOR signalling pathway affects different tissues in diverse ways. With many tissues, such as the heart, it is difficult to obtain sufficient biological material in amounts necessary for phosphoproteomics and interaction proteomics analyses of mTOR signalling in these tissues. Here we show that the first and crucial step to identifying tissue-specific mTOR substrates is to set up a tractable system to identify as many mTOR targets as possible. This is followed by identifying the substrates relevant to a specific tissue from the initial list identified through the proteomics approach^1^. We devised the SCGM strategy and successfully applied it to identify novel candidate mTOR substrates. We further demonstrate that the phosphorylation of LARP1, one of the identified mTOR substrates, is observed in the hearts of male mice with DCM, suggesting that LARP1 and the phosphorylation of LARP1 may be a biomarker for DCM.

A limitation of this strategy is that the quantitative and interaction proteomics analyses were performed in different cell lines and certain cell type specific interactors and substrates might not be identified. However, the high degree of homology of the mTOR protein sequences across human, mouse and rat, suggests that many interactors and substrates of mTOR may be conserved across these species and may be identified using this strategy.

The tractable aspect of our SCGM strategy means it can be applied to any cell line or cell type capable of producing sufficient cell numbers for proteomics analysis, such as cancer cell lines having abnormal mTOR signalling. mTOR inhibitors can be effective for treating certain cancers, including renal cell carcinoma^9,51^. However, cancer treatment with mTOR inhibitors may lead to resistance to these inhibitors^52^. Targeting cancer cell type-specific targets of mTOR for treatment would circumvent patients developing resistance to these inhibitors. Similarly, inhibition of activated mTOR in two mouse models of DCM significantly extends longevity^13,14^. Consequently, identifying mTOR substrates relevant to DCM progression would allow for the development of novel therapeutics targeting these cardiac-specific mTOR-substrate interactions, and as a result, potentially reduce the side-effects, such as immunosuppression, observed in patients treated with mTOR inhibitor, rapamycin^15^.

From our SCGM strategy, besides mTOR and some of its interactors/substrates, many of the proteins identified are not known to interact with mTOR. These novel candidate substrates participate in many biological processes, including responses to stressors, such as DNA damage and cell cycle arrest (Figure 2B). While mTOR signalling regulates these pathways to some extent, mTOR itself has not been shown to phosphorylate proteins comprising these pathways directly.

For instance, rapamycin, the allosteric inhibitor of mTOR, perturbs mTORC2 and alters cellular cytoskeletal organization^53,54^. However, the molecular pathway underlying the disruption of cytoskeletal organization is unknown. Here, we identified the microtubule-associated proteins, MAP1B and MAP4, as potential mTOR candidates. Phosphorylation of MAP1B and MAP4 regulates microtubule organization, and our finding that MAP1B and MAP4 are potential mTOR substrates provides a direct connection between mTOR and cytoskeletal organization^55,56^. Another biological pathway through which mTOR regulates and was identified in our study is rRNA processing. Previous studies showed that mTOR regulates rRNA processing indirectly through factors downstream of mTOR. UTP18 is a critical component of the processome required to correctly process pre-ribosomal RNA into the 18S ribosomal RNA^57^. The nucleo-cytoplasmic shuttling of UTP18 is rapamycin sensitive, suggesting that mTOR regulates the proper cellular localization of UTP18^57^. In our study, we identified UTP18 as a candidate mTOR substrate, again providing a direct link between mTOR and the regulation of rRNA processing. From these two examples, it is apparent that the data from our study is a valuable resource to aid in establishing the mode of action and pleiotropic functions of mTOR.

Intriguingly, several novel candidate mTOR substrates, namely, EDC4, RBM5, UTP18, SRPK2, HNRNPA1, DDX21, DDX24, and LARP1, are regulators of different aspects of RNA biology, ranging from pre-mRNA splicing to mRNA metabolism^34,57-63^. Identifying these proteins as potential mTOR substrates suggests that aside from regulating cellular protein homeostasis by governing mRNA translation, which mTOR is widely known for, mTOR also controls many facets of mRNA processing by directly phosphorylating these proteins. Studying the biological significance of the mTOR-dependent phosphorylation of these proteins would allow us to understand other means by which mTOR alters mRNA processing and metabolism to maintain cellular proteins levels in response to various cellular stimuli.

In two different *Lmna* mutant mouse lines that develop DCM, mTOR signalling levels were elevated in their hearts. Reducing mTOR signalling levels with mTOR inhibitors improved heart function in these affected mice by reducing ventricle size and increasing heart contractility^13,14^. Using another *Lmna*-mutant DCM mouse model *Lmna*^N195K^, we observed that, similar to the two previous studies, mTOR signalling levels increased in male mice with DCM compared to normal males^36^. Our results support the potential for inhibiting mTOR in treating DCM.

The mTOR-dependent phosphorylation of the substrate, EIF4EBP1, prevents EIF4EBP1 from inhibiting EIF4E (a protein translation initiator), promoting protein translation, including the translation of 5’ TOP mRNAs, a large portion of which encodes for ribosomal proteins^39^. However, several studies have demonstrated that while mTOR signalling regulates the translation of 5’ TOP mRNAs, the mTOR-dependent phosphorylation of EIF4EBP1 does not account for the selective regulation of 5’TOP mRNAs under different stimuli^32^. Our findings suggest that one of the stimuli-dependent regulation of proteins encoded by 5’ TOP mRNAs by mTOR is by regulating LARP1 phosphorylation.

Increased protein translation has been observed in premature ageing, and LARP1 regulates the stability of 5’ TOP mRNAs, including those translated into ribosomal proteins - proteins that are important for protein translation^33,64^. While the effect of mTOR phosphorylation of LARP1 on the translation of 5’ TOP mRNAs remain controversial, two recent studies demonstrated that the binding of LARP1 to 5’ TOP mRNAs stabilizes and promotes the translation of these mRNAs^33,34,68^. In line with these studies, our finding that mTOR-dependent phosphorylation of LARP1 enhanced the binding of LARP1 to 5’ TOP mRNAs suggests that this enhanced binding of phosphorylated LARP1 to RPS20 and RPL32 transcripts may stabilize these transcripts. The stabilization of these 5’ TOP mRNAs would, in turn, lead to more RPS20 and RPL32 proteins being translated and accumulated. The presence of elevated levels of RPS20 and RPL32 proteins observed in the hearts of male mice with DCM may be due to an increase in stability of these phosphorylated-LARP1 bound - 5’ TOP mRNAs^33^. In addition, our finding that the levels of ribosomal proteins are elevated in the hearts of mice with DCM suggests that aside from premature ageing, increased protein translation brought about by elevated levels of ribosomal proteins may be a significant contributor to the development of DCM^64^.

Further investigation into the identities of mRNA targets of LARP1 and how LARP1 translationally regulates them in cardiac cells will allow us to understand how elevated LARP1 and phospho-LARP1 levels may contribute to DCM progression. Recent studies on RNA binding proteins have revealed that these proteins represent attractive therapeutic targets for neurodegenerative disease and cancer^65,66^. Our finding that LARP1 might contribute to DCM suggests the possibility of targeting LARP1 for medical intervention for the treatment of DCM.

In summary, we have devised and employed our SCGM strategy to identify novel candidate mTOR substrates, many of which contribute to different aspects of RNA biology. This list of novel candidate mTOR substrates serves as an essential resource and starting point for studying tissue-specific mTOR interactions. Using the SCGM strategy, we established LARP1 as an mTOR substrate and determined that mTOR-dependent phosphorylation is nutrient-sensitive and alters the ability of LARP1 to bind its target mRNAs. In addition, our study is the first to provide a link between LARP1 and dilated cardiomyopathy.

## Supporting information

Tan et al. - Supplemental Data S4

Tan et al. - Supplemental Data S3

Tan et al. - Supplemental Data S2

Tan et al. - Supplemental Data S1

## Acknowledgements

We thank the Stewart and Burke lab members for their valuable discussions. We also thank Hui Jun Lim, Claire Tan and Gail Tan for critical reading of the manuscript, support and encouragement, and Vonny Ivon Leo for technical assistance. This research is funded by the Singapore Biomedical Research Council and the Singapore Agency for Science, Technology and Research (A*STAR) and a grant from the Progeria Research Foundation to CLS. RMS is supported by Core funding from IMCB and IMB Strategic Positioning Fund (SPF, BMRC, A*STAR), Young Investigator Grant YIG 2015 (BMRC, A*STAR), and NMRC MS-CETSA platform grant MOHIAFCAT2/004/2015

## Author Contributions

M-K.M.T and CLS designed the experiments and wrote the manuscript. M-K.M.T performed most of the experiments; RMS contributed to the mass spectrometry analyses; ESMW contributed to the mouse experiments; BB and LAV contributed to experimental design.

## Figure Legends

**Supplemental Figure 1:**
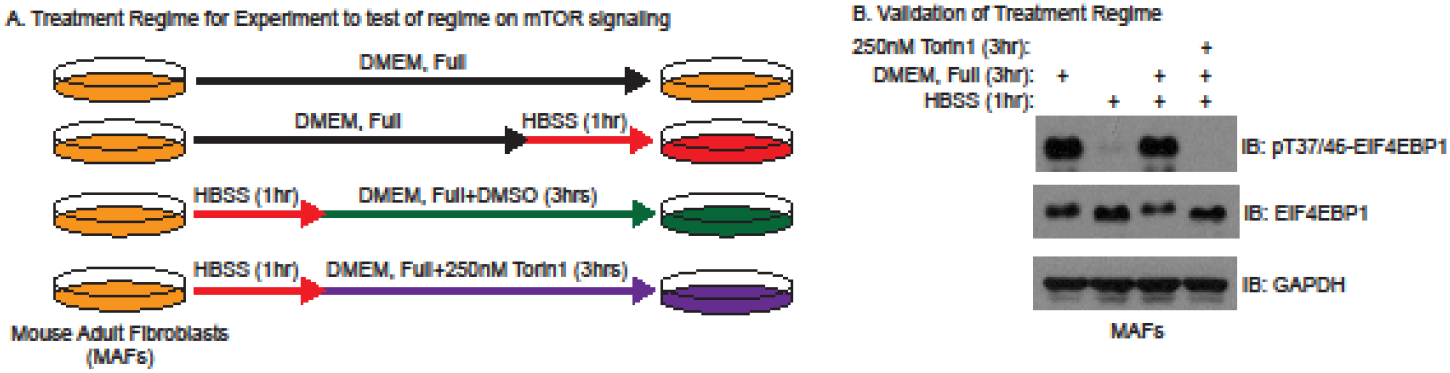
Testing of conditions for SILAC-phosphoproteomics. A) Schematic diagram of treatment regime using MAFs. B) Blot showing the result of immunoblotting with indicated antibodies after the appropriate treatments.

**Supplemental Figure 2:**
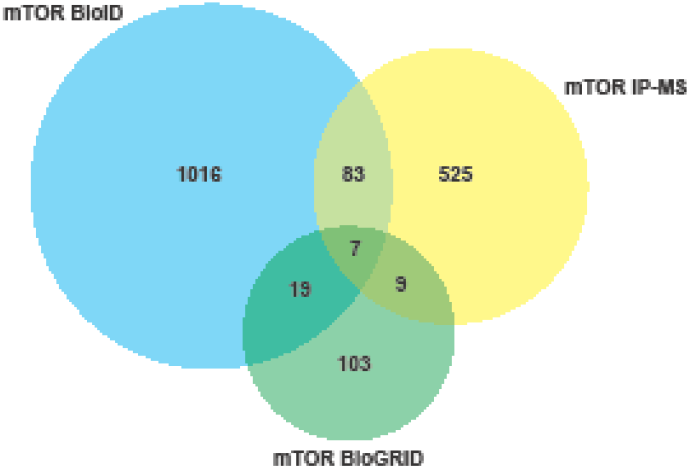
KEGG pathway analysis to determine common cellular pathways in which the proteins identified in our quantitative phosphoproteomics participate (p<0.05).

**Supplemental Figure 3:**
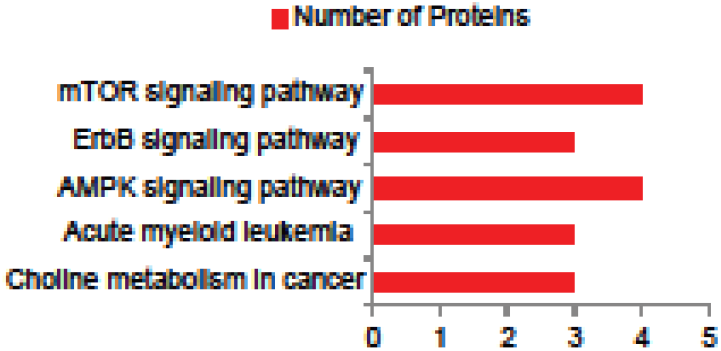
Venn diagram depicting the degree of overlap between the number of proteins found in each of our mTOR IP-MS and mTOR BioID studies with known mTOR interactions curated in BioGRID.

**Supplemental Figure 4:**
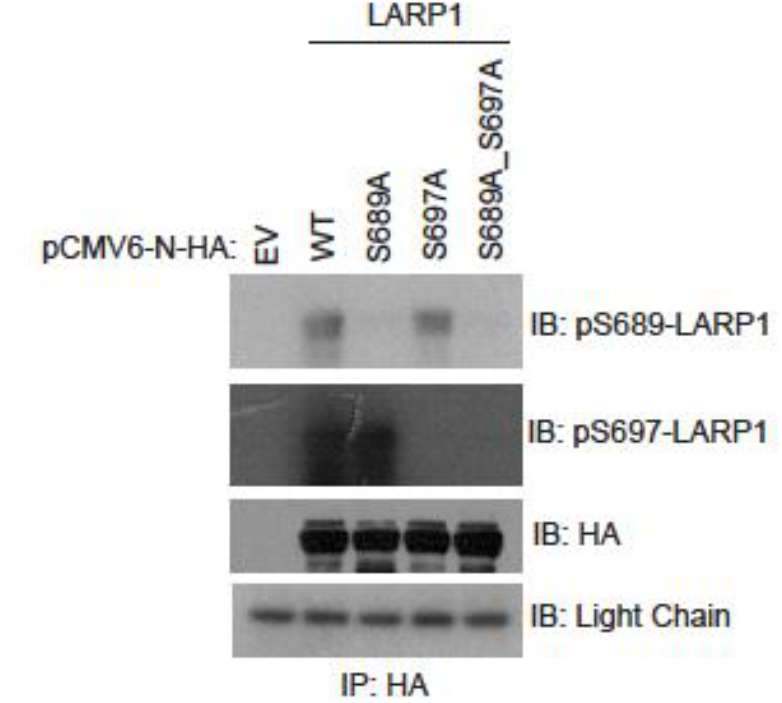
Testing of customized phospho-LARP1 antibodies. Immunoprecipitation using HA-conjugated agarose beads on the lysates of HEK293T cells transfected with the indicated HA plasmids was performed, followed by analysis with SDS-PAGE and immunoblotting with the indicated antibodies.

**Supplemental Data S1**: Tables showing normalized ratios of changes in peptide abundance between the compound-treated samples and the DMSO treated sample.

**Supplemental Data S2**: Table showing proteins with peptides with two or more fold changes in the compound-treated samples in both the Forward and Reverse Labelling experiments.

**Supplemental Data S3**: Tables showing proteins identified in the BioID and IP-MS experiments.

**Supplemental Table S4**: Table showing phosphorylated peptides of the 58 mTOR candidate substrates identified through the SCGM strategy. Phosphopeptides highlighted in red are peptides containing either a proline, phenylalanine or leucine exactly one position after the phosphorylated serine or threonine.

## Materials and Methods

### Cell culture and drug treatments

HEK293T and mouse adult fibroblasts (MAFs) cells were cultured in Dulbecco’s modified Eagle’s medium (DMEM) supplemented with 10% heat-inactivated fetal calf serum (FBS) and kept in a 5% CO2 incubator at 37°C. Rapamycin and Torin1 (both from Selleckchem) were resuspended in dimethyl sulfoxide (DMSO) (Sigma-Aldrich). The final concentrations of Rapamycin and Torin1 used in this study were 100nM and 250nM, respectively, unless otherwise stated.

### Stable isotope labelling by amino acids in cell culture (SILAC) of MAFs

MAFs were grown in stable isotope-labelled SILAC DMEM (Pierce) with 10% dialyzed FBS (Pierce) supplemented either with “Light” arginine and lysine (R0K0: R = 12C6, 14N4, K = 12C6, 14N2), “Medium” arginine and lysine (R6K4: R = 13C6, 14N4, K = 4,4,5,5-D4) or “Heavy” arginine and lysine (R10K8: R = 13C6, 15N4, K = 13C6, 15N2). All labelled amino acids were purchased from Cambridge Isotope Laboratories. All SILAC culture media was supplemented with L-proline (Sigma-Aldrich) to minimize arginine-to-proline conversion. These MAFs were grown for at least six cell doublings before they were treated with the appropriate compounds. SILAC amino acids incorporation was determined to be ∼95% after five cell doublings (Data not shown).

### Preparation of samples for Phosphoproteomics

In the Forward experiment, the SILAC samples were first nutrient-starved by replacing their growth medium with HBSS (Gibco) for 1 hour. After 45 minutes of HBSS incubation, DMSO, Rapamycin and Torin1 were added to the HBSS of the “Light”, “Medium”, and “Heavy” samples, respectively, for another 15 minutes. After the 1-hour HBSS incubation, the HBSS in the “Light”, “Medium”, and “Heavy” samples was removed and replenished with the corresponding SILAC growth media containing its respective treatment compound (either DMSO, Rapamycin or Torin1) and incubated for a further 3 hours. The process is similar for the Reverse experiment except in the Reverse experiment; the “Light”, “Medium”, and “Heavy” samples were treated with Rapamycin, Torin1 and DMSO, respectively. For both the Forward and Reverse Experiments, after the three hours incubation with the replenished SILAC growth medium containing the treatment compound, cells in the “Light”, “Medium” and “Heavy” samples were washed 3x with Phosphate Buffered Saline (PBS), dissociated with PBS-based enzyme-free cell dissociation buffer (Gibco), counted. Equal amounts of cells (∼3 × 107 cells) from each sample were pooled. The pooled cells were then lysed with Urea Lysis Buffer (8M urea, 75mM sodium chloride, 50mM Tris (pH 8.2), sonicated and centrifuged at maximum speed for 10 minutes at 4°C. The cleared lysate was transferred into a fresh tube, and its concentration was measured using the BCA protein assay (Pierce). The proteins in the lysate were reduced with 20mM Tris(2-carboxyethyl)phosphine (TCEP) (Sigma-Aldrich) and alkylated with 55mM 2-Chloroacetamide (CAA) (Sigma-Aldrich). The reduced and alkylated proteins in the lysate were first digested overnight with Lysyl Endopeptidase (LysC) (Wako), followed by another overnight digestion with sequencing grade Trypsin (Pierce). Trypsin digestion was stopped by the addition of Trifluoroacetic acid (TFA) (Sigma-Aldrich) to a final concentration of 0.4%, and the digested peptides obtained were desalted using a Sep-Pak C18 Classic Cartridge (Waters). The cartridge was conditioned by first washing with 9mL acetonitrile (ACN), followed by 3mL elution buffer (50% ACN and 0.5% acetic acid (HAcO)) and a subsequent wash with 9 ml 0.1% TFA solution. After conditioning the cartridge, the sample was loaded, and the flow-through was then reloaded into the cartridge. The cartridge was washed with 900mL 0.5% HAcO solution and eluted with 2ml elution buffer. The elution was performed another two times by reloading the eluate into the cartridge. The eluate obtained was lyophilized and stored at 80 °C.

### Phosphopeptide enrichment

Enrichment was performed using SCX combined with TiO2 method essentially described in Olsen, J.V. et al., (2006), Cell^27^. Prior enrichment input control was collected. The remaining samples were vacuum centrifuged and diluted with SCX solvent A (10mM KH2PO4, 20%ACN pH 2.7) further subjected to separation in the 60min salt gradient up to 35% of Solvent B (350mM KH2PO4, 20% ACN) using SCX Resource S column and AKTA Micro liquid chromatography system (GE). Collected fractions were combined and subjected to TiO2 batch enrichment (16 fractions).

### Mass Spectrometry

Analysis was performed using Easy nLC1000 (Thermo) chromatography system coupled with Orbitrap Fusion (Thermo). Each sample was separated in 120min gradient (0.1% Formic Acid in water and 99.9% Acetonitrile with 0.1% Formic Acid) using 50cm x 75um ID Easy-Spray column (C-18, 2um particles, Thermo). Data-dependent mode for phosho proteomic analysis was used in a speed mode -3 sec cycle using Orbitrap and ion trap analyzers (OT-MS 4xE5 ions, resolution 60K, IT MS/MS 6000 and HCD fragmentation).

### Data processing

Peak lists were generated with MaxQuant software^67^. Searches were done against forward/decoy Mouse Uniprot database with the following parameters: precursor mass tolerance (MS) 20ppm, IT-MS/MS 0.5 Da, two miss cleavages. Static modifications: Carbamidomethyl (C), SILAC amino acids Arg+6, Lys +4, Arg+10, Lys +8. Variable modifications: Oxidation (M), Deamidated (NQ), Phospho (STY), Acetyl (N-terminal protein). Forward/decoy searches were used for false discovery rate estimation (FDR 1%). Following analysis, the input sample was used for phosphopeptides normalization.

### Preparation of BioID samples

HEK293T cells (∼107) were transiently transfected with either pcDNA3.1-N-MYC-BioID2-EGFP (control), pcDNA3.1-N-MYC-BioID2-human mTOR (hmTOR), pcDNA3.1-N-MYC-BioID2-rat mTOR (rmTOR) using the TransIT-X2 transfection reagent (Mirus) according to the manufacturer’s specification. Thirty hours post-transfection, the transiently transfected HEK293T cells were treated with 50uM Biotin (Sigma-Aldrich) for 12 hours. After 12 hours, the HEK293T cells were starved by replacing the growth media with HBSS for an hour. After an hour in HBSS, growth media was replenished to all the samples, and the cells were grown in fresh media for another 3-4 hours. The HBSS and replenished growth media both contained 50uM Biotin. The samples were subsequently harvested and washed three times with PBS before being lysed with BioID cell lysis buffer (50mM Tris pH 7.4, 500mM NaCl, 0.4% SDS, 5mM EDTA, and 1mM DTT, 2% Triton-X 100, Protease and Phosphatase inhibitors (Roche)). After lysis, the lysate obtained for each sample was sonicated on ice, followed by centrifugation in the cold at maximum speed. 100uL of equilibrated Streptavidin-coupled Dynabeads (ThermoFisher) were added to each sample, and the samples were left to rotate overnight at 4°C. The next day, the Dynabeads were separated from the supernatant by placing the tubes on a magnetic holder and aspirating the supernatant. The recovered Dynabeads were first washed twice with Wash Buffer (WB) 1 (1% SDS in water), followed by a wash with WB2 (0.1% sodium deoxycholate, 1% Triton X-100, 500mM NaCl, 1mM EDTA), a wash with WB3 (250mM LiCl, 0.5% NP-40, 0.5% sodium deoxycholate, 1mM EDTA, 10mM Tris pH 8.0) and lastly with two washes with WB4 (50mM Tris pH 7.4, 50mM sodium chloride). WB4 was removed from the beads, and the recovered beads were used for LysC and Trypsin digestion.

### Preparation of Immunoprecipitation (IP) samples for mass spectrometry analysis

HEK293T cells were transiently transfected as those in the BioID analysis. Forty hours post-transfection, the HEK293T cells were starved by replacing the growth media with HBSS for an hour. After an hour in HBSS, growth media was replenished to all the samples, and the cells were grown in fresh media for another 3-4 hours. The samples were then harvested and washed twice with PBS before being lysed with 0.3% CHAPS lysis buffer (40 mM HEPES, pH7.5, 0.3% [w/v] CHAPS, 120mM sodium chloride, 1mM EDTA, 0.5mM DTT, 1ug/mL RNase, Protease and Phosphatase Inhibitors (Roche)). The lysate from each sample was cleared by centrifugation at maximum speed in the cold. 50uL of equilibrated MYC antibody-conjugated Dynabeads (ThermoFisher) were added to each of the cleared lysates, and the tubes containing the lysate and beads were rotated overnight in the cold. The next day, the Dynabeads were separated from the supernatant by placing the tubes on a magnetic holder and aspirating the supernatant. This was followed by washing the beads five times with 0.3% CHAPS lysis buffer. The washed beads were recovered and used for LysC and Trypsin Digest.

### LysC and Trypsin Digest, and peptide desalting

The beads in each sample from the BioID and IP proteomics experiments were resuspended in buffer containing 50mM Tetraethylammonium borohydride (TEAB) (Sigma-Aldrich-Aldrich) and 50% Trifluoroethanol (TFE). After resuspension, each sample was reduced with 20mM TCEP and alkylated with 55mM chloroacetamide (CAA) (Sigma-Aldrich). The reduced and alkylated samples were then digested with 10ug LysC (Wako) and 10ug Trypsin (Pierce). After the digest, the tubes containing the samples were placed on a magnetic rack to separate the beads from the supernatant. The supernatant containing peptides from each sample was transferred to a new 1.5mL tube. Trifluoroacetic acid was added to each sample to a final concentration of 1% (v/v). Each sample was then desalted with equilibrated Sep-Pak C18 Classic Cartridge (Waters) and was passed through a C18 cartridge, followed by washing the cartridge with buffer containing 0.5% acetic acid (Sigma-Aldrich-Aldrich). The desalted peptides in the cartridge were eluted using a buffer containing 0.5% acetic acid and 80% acetonitrile (Sigma-Aldrich-Aldrich). The samples were subsequently dried using a speedvac for mass spectrometry analysis.

### Antibodies

The following antibodies were used: α-RPS20 (ab133776; Abcam), α-RPL32 (ab50759; Abcam), α-LARP1 (ab86359; Abcam, 13708-1-AP; Proteintech), α-PABP (ab21060; Abcam), α-EIF4A1 (ab31217; Abcam), α-RAPTOR (sc-81537; Santa Cruz), α-SRPK2 (sc-390534; Santa Cruz), α-DTL (A300-948A-T; Santa Cruz), α-GAPDH (ab8245; Abcam), α-pT37/46-4E-BP1 (2855S; Cell Signalling Technology), α-pT70-4E-BP1 (13396S; Cell Signalling Technology), α-4E-BP1 (9644S; Cell Signalling Technology), α-pS2488-mTOR (2971S; Cell Signalling Technology), α-mTOR (2972S; Cell Signalling Technology), α-HA (MMS-101P; Covance), α-Myc-Tag (2276; Cell Signalling Technology), α-DDX21 (A300-628A; Bethyl), both α-pS689-LARP1 and α-pS697-LARP1 were manufactured by YenZym Antibodies LLC.

### Cell Lysis and Immunoprecipitation (IP)

Cells were lysed either with 1% NP-40 lysis buffer [50 mM Tris-HCl (pH 7.5), 150 mM NaCl, 1% NP-40, 1 mM EDTA and supplemented with protease and phosphatase inhibitors (Roche)] or with 0.3% CHAPS lysis buffer [40 mM HEPES, pH7.5, 0.3% [w/v] CHAPS, 120 mM sodium chloride, 1 mM EDTA, 0.5mM DTT, 1ug/mL RNase, protease and phosphatase inhibitors (Roche)] for 20 min on ice. The lysate obtained from each sample were cleared by centrifugation at maximum speed at four °C for 15 min. For endogenous IP, approximately 2 mg of lysate and 25uL of Protein G Dynabeads (ThermoFisher) were incubated with 1 μg of the indicated antibody or control IgG overnight at 4°C. The beads were recovered and washed three times with the appropriate lysis buffer. After washing, the beads were resuspended in 2x SDS loading buffer and boiled for 5 min. Samples were separated on an SDS-PAGE gel before immunoblot analysis. For IPs concerning ectopically expressed proteins, at least 400ug of lysate and 25uL of Dynabeads conjugated with either HA-antibody (ThermoFisher) or MYC-antibody (ThermoFisher), or agarose beads conjugated with either HA antibody (Sigma-Aldrich) or MYC antibody (Sigma-Aldrich) was used for each sample.

### Lambda Phosphatase Treatment

Three sets of HEK293T cells were first starved by growing them in HBSS for one hour. After an hour, HBSS was removed from all three groups of cells. Two groups of cells were then cultured for another 3 hours in complete growth media with DMSO, while another set of cells were cultured for 3 hours with complete growth media with 250nM Torin1. After the three-hour growth media incubation, the cells were harvested and lysed with 1% NP-40 lysis buffer [50 mM Tris-HCl (pH 7.5), 150 mM NaCl, 1% NP-40, one mM EDTA and supplemented with protease and phosphatase inhibitors (Roche)]. One set of cells cultured in complete growth media with DMSO were lysed with 1% NP-40 lysis buffer without phosphatase inhibitor. The lysate obtained from lysis with 1% NP-40 lysis buffer phosphatase inhibitor were mixed with lambda phosphatase (New England Biolabs) and incubated at 30°C for 2 hours. After the lambda phosphatase treatment, 2x SDS loading buffer was added to the lysates obtained from all three sets. The samples were then boiled for 5 min. Samples were separated using SuperSep Phos-tagTM acrylamide gel (Wako) and processed according to the manufacturer’s protocol before immunoblotting.

### In vitro kinase assays

HEK293T cells (∼107) were transiently transfected with either pRK5-MYC-mTOR (Addgene plasmid #1861) or pRK5-MYC-mTOR kinase-dead (Addgene plasmid #8482)) using the TransIT-X2 transfection reagent (Mirus) according to the manufacturer’s specification. Both the pRK5-MYC-mTOR and pRK5-MYC-mTOR kinase-dead plasmids were gifts from David Sabatini. Forty hours post-transfection, 10ug/mL insulin (Sigma-Aldrich-Aldrich) were added to the transfected cells for 30 minutes. After 30 minutes, the cells were washed once with phosphate-buffered saline and lysed with 0.3% CHAPS lysis buffer [40 mM HEPES, pH7.5, 0.3% [w/v] CHAPS, 120 mM sodium chloride, 1mM EDTA, 0.5mM DTT, 1ug/mL RNase, protease and phosphatase inhibitors (Roche)] on ice for 30 minutes. The lysed cells were then centrifuged at maximum speed at 4°C for 15 minutes. The cleared lysates were transferred to fresh 1.5mL tubes and incubated with 0.3% CHAPS lysis buffer and MYC-conjugated agarose beads (Sigma-Aldrich-Aldrich) overnight at 4°C. After the overnight incubation, the beads were first washed with low salt mTOR wash buffer containing 40 mM HEPES (pH 7.4), 150 mM NaCl, 2mM EDTA, protease and phosphatase inhibitors (Roche) with 0.3% CHAPS (v/v). They were then washed twice with high salt mTOR wash buffer containing 40 mM HEPES (pH 7.4), 400 mM NaCl, 2mM EDTA, protease and phosphatase inhibitors (Roche) with 0.3% CHAPS (v/v), followed by another two washes with mTOR wash buffer containing 25 mM HEPES (pH 7.4), 20 mM KCl. The washed beads were divided equally into three 1.5mL tubes.

To obtain the bacterial purified GST-EIF4EBP1, GST-LARP1-WT (417-1019) and GST-LARP1-DM (417-1019) proteins, cDNA encoding either EIF4EBP1, LARP1-WT (417-1019) or LARP1-DM (417-1019) were cloned into the pGEX-6P1 vector. After cloning, the plasmids were used to transform BL21 (DE3) chemically competent cells (Sigma-Aldrich-Aldrich). After transformation and growing the transformed cells on antibiotics selection agar plates, a single colony for each sample was gown in LB bacterial growth media until an OD590 of 06-08. After that, isopropyl-β D-thiogalactoside (IPTG) was added to each sample to give a final concentration of 0.4mM and incubated for 5h at 37°C. The cells were then pelleted, resuspended in PBS supplemented with 0.1mg/mL lysozyme, and incubated on ice for 30 minutes. This was followed by pulse sonication to shear bacterial DNA. After sonication, the samples were centrifuged at 16,000xg for 20 minutes at 4°C. The supernatants from each sample were then incubated with glutathione-sepharose beads (GE Healthcare Life Sciences) to immunoprecipitate the GST tagged proteins. After immunoprecipitation, the beads were washed and incubated with 50U shrimp alkaline phosphatase (New England Biolabs) for 30 min at 37°C. The beads were then washed with PBS supplemented with 10mM EDTA and 0.1% (v/v) Triton X-100. The GST-tagged proteins were eluted from the beads using 50mM Tris-HCl, 10mM reduced glutathione pH=8. Buffer exchange was performed using Amicon Ultra-15 centrifugal filter units (Merck) to have the purified proteins in mTOR wash buffer containing 25 mM HEPES (pH 7.4), 20 mM KCl.

The kinase assay was performed by adding 150ng of bacterial purified GST proteins to one tube each of the MYC-mTOR (WT) and MYC-mTOR kinase-dead (KD) immunoprecipitates in mTORC1 kinase buffer containing 500uM ATP, 25mM HEPES (pH 7.4), 50mM KCl, 10mM MgCl2 and Incubated at 30°C for 60 minutes with shaking. The reactions were stopped by adding 5x SDS sample buffer. Subsequently, the samples were analyzed by first running them in an SDS-PAGE gel followed by immunoblotting.

### RNA immunoprecipitation and quantitative PCR (qPCR)

HEK293T cells (∼107) were transiently transfected with either pCMV6-AN-HA-EGFP (control), pCMV6-AN-HA-LARP1-WT, pCMV6-AN-HA-LARP1-DM using the TransIT-X2 transfection reagent (Mirus) according to the manufacturer’s specification. Forty-eight hours post-transfection, the cells were washed once with 10mL phosphate-buffered saline, harvested and lysed with polysome lysis buffer containing 100mM KCl, 5mM MgCl2, 10mM HEPES, pH 7.0, 0.5% Nonidet P-40, 1mM dithiothreitol (DTT), 200 units/ml RNase Inhibitor, one Complete Mini, EDTA-free Protease Inhibitor Tablet (Roche) on ice for 30 minutes. After lysis, the lysates were centrifuged at 20,000xg for 10 minutes at 4°C. The cleared lysates were transferred to fresh 1.5mL tubes, and 100uL of lysate from each sample were added to 800uL of NET-2 buffer containing 50mM Tris pH 7.4, 150mM NaCl, 1mM MgCl2, 0.05% Nonidet P-40, 20mM EDTA pH 8.0, 1mM DTT, 200 units/ml RNase Inhibitor, and 100uL of HA-antibody conjugated agarose bead slurry. The lysate and HA-antibody conjugated agarose bead mixtures were centrifuged briefly to allow the beads to settle at the bottom of the tubes. 100uL of the lysate and NET-2 mixture above the beads was collected in a new tube and labelled ‘total’. The remaining lysate, NET-2 and beads mixtures were incubated overnight at 4°C. After the overnight incubation, the beads were washed six times with ice-cold NT-2 buffer containing 50mM Tris pH 7.4, 150mM NaCl, 1mM MgCl2, 0.05% Nonidet P-40. The washed beads and the ‘total’ samples were then resuspended in Proteinase K buffer containing 50mM Tris pH 7.4, 150mM NaCl, 1mM MgCl2, 0.05% Nonidet P-40, 1% sodium dodecyl sulfate (SDS), 1.2 mg/ml Proteinase K, and incubated at 55°C for 30 minutes.

After the Proteinase K treatment, RNA was extracted from the samples using TRIzol® according to the manufacturer’s protocol. 1ug of total RNA from each sample was used for reverse transcription using the High Capacity cDNA Reverse Transcription Kit (Applied Biosystems) according to the manufacturer’s protocol. cDNA obtained was used for qPCR analysis using TaqmanTM (ThermoFisher) probes for human RPS20 and RPL32 (Assay IDs Hs00828752_gH and Hs00851655_g1, respectively) according to manufacturer’s protocol.

### Animal Work

Mouse genotyping was performed by PCR. All animal experiments were performed following the BRC Animal Use Committee approved animal study protocols.

